# Eco-evolutionary dynamics between multiple competitors reduce phytoplankton coexistence but have limited impacts on community productivity

**DOI:** 10.1101/2025.01.13.632708

**Authors:** Charlotte L. Briddon, Aurora Menéndez García, Giulia Ghedini

**Affiliations:** Instituto Gulbenkian de Ciência, Rua Quinta Grande 6, 2780-156, Oeiras, Portugal; Ghent University, Department of Biology, Marine Biology Research Unit, Krijgslaan 281/S8, 9000 Ghent, Belgium; School of Biological Sciences, Monash University, Clayton, VIC 3800, Australia

**Keywords:** metabolism, community functioning, biomass, biodiversity, evolution, competition

## Abstract

Species can evolve rapidly in response to competition but how evolution within communities affects community properties is unclear. To test this, we grew three marine phytoplankton species in monoculture (alone) or polyculture (together) for 17 weeks. We then combined them in communities based on their competition history (monoculture or polyculture isolates) and tracked their composition and productivity over time. We found that species dominance was unaffected, but coexistence was reduced when species evolved together (polyculture isolates). Total biovolume was robust to changes in species relative abundances. However, polyculture isolates had greater oxygen fluxes during exponential phase and were less robust to the addition of an invader. Our results suggest that evolution within communities can strengthen competitive differences between species with uneven effects on different aspects of community functioning. Thus, we should be cautious in extrapolating the consequences of evolution on community biomass to other aspects of productivity or stability.

## INTRODUCTION

Ecological interactions can result in rapid evolution. While there is increasing evidence of the effects of eco-evolutionary dynamics on the traits of organisms (Hart *et al*. 2019; Terhorst *et al*. 2018), the consequences on community properties remain unclear, particularly for eukaryotes. In the presence of competitors organisms often evolve at different rates or in different directions than organisms alone (Lawrence *et al*. 2012; De Mazancourt *et al*. 2008). Understanding if communities of species that evolved together have different properties than those composed of novel species is particularly important given that biodiversity changes are increasing the probability that species will encounter new competitors.

Theory predicts that species that compete for similar resources should evolve distinct phenotypes when they coexist in the same areas (sympatry) compared to when they exist in isolation (Pinsky 2019; Scheuerl *et al*. 2020; Veresoglou *et al*. 2024). Such character divergence can facilitate coexistence by promoting niche partitioning (Gibbs *et al*. 2022; Lear *et al*. 2021; Pfennig & Pfennig 2009). Patterns of niche differentiation are often observed in bacteria where species can evolve to preferentially use the waste products of other species, thus increasing facilitative interactions (Lawrence *et al*. 2012), although that is not always the case (Castledine *et al*. 2020). Over generations, trait divergence can improve species fitness and increase community productivity due to more efficient resource use (Lawrence *et al*. 2012; Piccardi *et al*. 2022).

In species that compete for essential resources, such as plants or algae, the evolution of niche partitioning might be rare (Abrams 1986). In these cases, competition might more frequently alter competitive ability rather than the type of resources used (Gorter *et al*. 2020; Sakarchi & Germain 2023). Changes in competitive ability might have important consequences for community structure and functioning. For instance, changes in competitive ability might facilitate coexistence if they occur in parallel across species because they can reduce fitness differences (Hart *et al*. 2019). Improvements in competitive ability can also enhance the overall efficiency of resource use (Ghedini *et al*. 2020; MacArthur 1969) potentially constraining the establishment of invaders (Germain *et al*. 2020). However, beyond bacteria (Castledine *et al*. 2020; Lawrence *et al*. 2012), tests for how evolutionary processes between multiple competitors affect community properties are rare (De Meester *et al*. 2019).

Phytoplankton provides a useful model system to study evolution in communities of eukaryotic organisms because species have rapid cell division (∼1 per day; Banse 1991; Cooper 2000), large population sizes and therefore high capacity for evolution (Collins *et al*. 2014; Padfield *et al*. 2016). These organisms perform important ecosystem functions because they transfer energy to foodwebs and drive the carbon cycle (Litchman *et al*. 2007). So, understanding how eco-evolutionary processes influence the functioning of phytoplankton is particularly important. However, maintaining multiple competitors in the same environment for enough time to allow evolution is difficult in the laboratory (Vanvelk *et al*. 2024). To solve this challenge, we used an experimental approach which is a close but imperfect representation of evolution in a community. Specifically, we experimentally evolved three phytoplankton species together in a community (polyculture) for 4.5 months by enclosing each species in a separate “cage” (dialysis bag) – so that species competed for nutrients but were physically separated (Figure 1). We did the same thing for species alone (enclosing replicate populations of the same species in dialysis bags). Dialysis bags have been used in many experiments before to expose organisms to the same environmental conditions while physically keeping them separated (Ghedini & Marshall 2023; Pomati *et al*. 2017; Poulson-Ellestad *et al*. 2014; Reed & Martiny 2013; Scheuerl *et al*. 2020). When we phenotyped these species individually, after 4.5 months of evolution, we found that they all increased their competitive ability by reducing their sensitivity to intraspecific competition (lower density-dependence of growth and energy use), but these responses were weaker for species in polyculture (Briddon *et al*. 2024).

**Figure 1.**
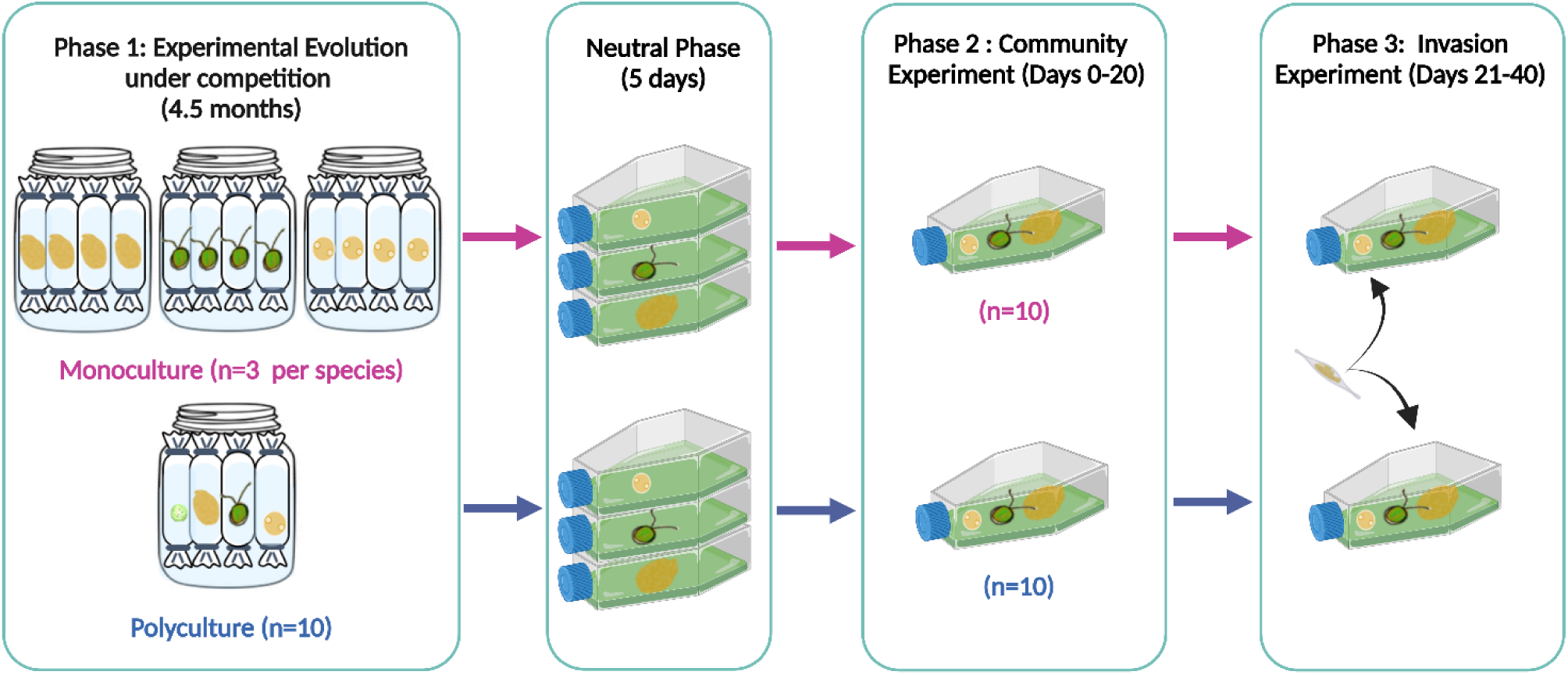
Schematic showing the different experimental phases. During Phase 1 (Experimental Evolution), we evolved three phytoplankton species (*Amphidinium*, *Dunaliella*, *Tisochrysis*) either under intraspecific (monoculture) or interspecific competition (polyculture) for 4.5 months, using dialysis bags that allow competition but keep these species separate (the fourth species, *Nannochloropsis*, was competitively excluded halfway during the evolution phase and thus was not included in the following experiments). After 4.5 months, we combined the species with a shared competition history for Phase 2 (Community experiment; Days 0-20) and Phase 3 (Invasion; Days 21-40). Our goal was to assess how competition history impacted community functioning and its stability to an invader. Before Phase 2, we grew each species in an individual flask for 5 days to remove any plastic response (neutral phase).

Our goal here is to determine the community effects of these evolved changes and whether they differ for communities of species evolved alone (monoculture isolates) or together (polyculture isolates). We hypothesise that 1) polyculture isolates should show greater coexistence than monoculture isolates because species had the opportunity to adapt to interspecific competitors. While phytoplankton compete for essential resources, these species can partition some resources such as light (Heggerud *et al*. 2023; Stomp *et al*. 2004) so evolution with competitors might result in (some degree of) niche differentiation. Furthermore, all species reduced their sensitivity to intraspecific competition, thus potentially equalising fitness differences. We also predict that 2) polyculture isolates should be more productive and resistant to invasion because resources should be more fully utilised by the community members (either through niche partitioning or increased resource use efficiency), preventing the establishment of an invader.

## MATERIALS AND METHODS

To test the consequences of evolution with competitors on community assembly and functioning, we used three species of marine phytoplankton (*Amphidinium carterae* RCC88*, Dunaliella tertiolecta* RCC6, and *Tisochrysis lutea* RCC90) acquired from the Roscoff Culture Collection, France. These species belong to different taxonomic groups and have different morphologies and cell sizes, encompassing different competitive and physiological traits (*Amphidinium;* initial average cell size Monoculture isolates = 779.53 ± 32.14 µm^3^; Polyculture isolates = 773.07 ± 16.29 µm^3^; *Dunaliella* = 387.03 ± 28.52 µm^3^; 375.86 ± 15.13 µm^3^; *Tisochrysis =* 71.53 ± 3.34 µm^3^; 70.46 ± 1.69 µm^3^). First, we evolved these species either alone (“monoculture”) or together (“polyculture”) for 4.5 months (Phase 1: Evolution experiment ∼120 generations). The evolution phase also included a fourth species, *Nannochloropsis granulata* (19 ± 0.17 µm^3^; RCC438), from the same phytoplankton collection. However, *Nannochloropsis* was competitively excluded in the polyculture treatment after 16 weeks, therefore it was not included in subsequent experiments. After Phase 1, we combined the evolved species in communities based on their competition history to determine how communities of species evolved together (polyculture isolates) perform relative to novel communities (monoculture isolates) (Phase 2: community experiment). Once the communities approached carrying capacity, we introduced an invader, the diatom *Phaeodactylum tricornutum* (RCC69; 69.41 ± 26.01 µm^3^), to test the stability of communities to a species that was never encountered (Phase 3: invasion).

### Phase 1: Experimental Evolution

To evolve each species in monoculture or polyculture we used dialysis bags (MWCO 14 kDa, pore size 25 Angstrom, Dialysis membrane Membra-CEL, Carl Roth, Germany), which allowed competition for nutrients and exchange of metabolites but maintained a physical separation between species (Ghedini & Marshall 2023; Pomati *et al*. 2017; Rotem *et al*. 2010). For the intraspecific treatment (monoculture), we established three replicate beakers per species. Each beaker contained four dialysis bags, each filled with the same species surrounded by enriched seawater medium (f/2 media prepared from 0.2 µm filtered and autoclaved natural seawater (Guillard & Ryther 1962)) containing no phytoplankton. For the interspecific treatment (polyculture), we set up 10 replicate beakers using the same design but each of the four dialysis bags within a beaker was filled with a different species (Figure 1). Once a week, we transferred a set volume of 25 ml from each dialysis bag to a new sterilised bag and beaker. The contents of the bag were topped up with 20 ml of fresh media. We also replaced the media in the beaker. Light intensity was set at 60 µmol m^-2^ s^-1^ with a 12:12 light:dark schedule using low-heat 50W led flood lights (Surface Luminária LED 230V, Robert Mauser, Portugal). The temperature was maintained at 20 ± 1°C.

### Phase 2: Community Experiment

After 4.5 months of evolution, we collected 10 samples (20 ml) for each species and competition treatment (monoculture and polyculture, N = 60) and placed them alone in a neutral environment (cell culture flasks filled up to 100 ml with f/2 media) for five days (∼ 5 generations) to remove any environmental conditioning. For the monoculture treatment, we sampled multiple bags (populations) from the same beaker (since we had 12 bags in total divided between 3 beakers per species). We recognise this is not ideal because these populations are not independent replicates as they shared the same beaker. However, we did not find “beaker” effects on the traits of the individual populations (Briddon *et al*. 2024); furthermore, the populations evolved separately within each bag for 4.5 months, so they do not share genetic mutations that appeared during this time.

After the neutral phase, we used the populations in the individual flasks to assemble communities of novel species (by randomly combining monoculture isolates of the three species) and communities of species that evolved together (by combining polyculture isolates, in this case we combined species that came from the same beaker; n = 10 for each community type). All communities were started with an equal biovolume of each species (1.06 × 10^4^ µm^3^ µl^-1^) which was added to 250 ml culture flasks, filled up to 100 ml with f/2 media (Figure 1). Phase 2 lasted for 20 days, until the samples reached carrying capacity. Each sampling day, we collected 10 ml from each culture flask for analysis (see details below) and replaced it with 10 ml of fresh media. The sampling took place on days 2, 4, 6, 8, 10, 12, 14, 17 and 20.

### Phase 3: Invasion

Once the communities reached stationary phase (∼ day 20), we introduced (on day 22) the invader species (*Phaeodactylum tricornutum)*, using the same initial biovolume used for the other species (1.06 × 10⁴ µm³ µl⁻¹). The sampling procedures were identical to those used in Phase 2 (the community experiment) with measurements taken on days 23, 26, 28, 30, 33, 34, 37, and 40.

### Cell size, densities and biovolume

On each sampling day during Phase 2 and 3, we measured cell size (µm^3^), cell circularity, cell density (cells/µl) and biovolume (µm^3^/µl; calculated by multiplying cell density and average cell size for each replicate). We followed the procedure detailed in Briddon *et al*. (2024). Briefly, we fixed 1 ml of sample from each replicate with 1% Lugol and acquired 20 photos with an inverted Olympus microscope (400x magnification). The size and abundance of each species was recorded using ImageJ and Fiji software (version 2.0) (Schindelin *et al*. 2012).

### Metabolic Rates

In conjunction with biovolume data, we measured photosynthesis, post-illumination and dark respiration rates to determine changes in community metabolism. For each community, we collected 5 ml and placed it in a vial with an integrated oxygen sensor. Vials were placed on 24-channel optical fluorescence oxygen readers to measure oxygen production and consumption (PreSens Sensor Dish Reader, SDR; PreSens Precision Sensing, Germany) (Briddon *et al*. 2024; Ghedini & Marshall 2023). The samples were measured for 20 minutes in the light to quantify photosynthesis (the first three minutes of measurements were discarded to allow acclimation), followed by 40 minutes of darkness to determine respiration rates (the first 15 minutes were used to quantify the post-illumination rates and the remainder the dark respiration rates). As the cultures were not axenic, eight blanks per day were filled with spent media with no phytoplankton cells to account for the background bacterial activity (spent media was obtained by centrifuging samples at 5,000 rpm for 10 minutes to separate the algal cells from the supernatant).

The photosynthesis and respiration rates (VO_2_; μmol O_2_/min) of each sample were calculated as:

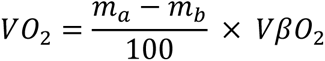

where m_a_ is the rate of change in O_2_ percentage of the sample (min^-1^), m_b_ is the mean O_2_ % across all blank samples (min^-1^), V is the sample volume (0.005 L) and βO_2_ is the O_2_ capacity of air-saturated seawater at 20°C and 35 ppt salinity (225 µmol O_2_/L) (White *et al*. 2011). Subsequently, we converted the photosynthetic and respiration rates (µmol O_2_/min) to calorific energy (J/min), using a conversion factor of 0.512 J/µmol O_2_ to estimate the energy production and consumption respectively (Williams & Laurens 2010). These calculations were completed using the LoLinR package (Olito *et al*. 2017) in R (version 4.3.2) (R Core Team 2021).

### Data Analysis

The statistical analyses were done on the data collected during Phase 2 (Community Experiment) and Phase 3 (Invasion). All data were analysed using R (version 4.3.2) (R Core Team 2021) and Rstudio (RStudio Team 2021) with the packages nlme (Pinheiro 2022), lme4 (Bates *et al*. 2015), emmeans (Searle & Milliken 1980), car (Fox 2019), plyr (Wickham 2011b), vegan (Oksanen *et al*. 2024) for analyses and ggplot2 (Wickham 2011a; Wilke 2016). For all statistical analyses, any interactions between the covariates were removed when p > 0.25. All figures with means show the least square means with a 95% confidence interval using a Tukey p-value adjustment unless stated otherwise.

### a. Differences in species growth dynamics

Using data from Phase 2, we tested how each species performed in the community based on its competition history (monoculture, polyculture). Using biovolume data (μm^3^/μl), we calculated the maximum rate of increase (r_max_) and the maximum value (i.e. carrying capacity, K) for each species within each community using the same approach described in Briddon *et al*. (2024). Briefly, we fit four models to biovolume data: a logistic-type sinusoidal growth model with lower asymptote forced to 0, a logistic-type sinusoidal growth model with non-zero lower asymptote, a Gompertz-type sinusoidal growth model and a modified Gompertz-type sinusoidal growth model including population decline after reaching a maximum. Then we used AIC (Akaike information criterion) to determine the best fitting model for each population with successful convergence. From this model, we estimated the maximum predicted value (K) of biovolume for each species in each community. From the first derivative, we extracted the maximum rate of increase (r_max_). We then tested for differences in these parameters (r_max_ and K) of each species separately using a linear model that included initial biovolume as covariate and competition history (monoculture and polyculture isolates) as factor, including their interaction.

To visualise differences in the growth trajectory of each species between the two community types, we used the same models above, but we fitted them across all replicates of each species within a treatment. We then plotted the best-fitting model based on AIC.

### b. Changes in cell size

For each species (Phase 2), we tested for differences in cell size and shape due to their competition history. We used a linear mixed effect model that included competition history and experiment day as factors (with their interactions), with sample code as a random effect to account for repeated measures.

### c. Evenness

We calculated Pielou’s evenness index (J’) based on the biovolume of the individual species to estimate changes in community composition (including species diversity and relative abundances) using the data from Phase 2 (Smith & Wilson 1996):

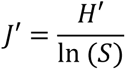

where H’ Shannon-Weiner diversity index and S is the number of species in the sample. To determine if species evenness differed due to competition history, we used a linear mixed effect model with evenness as a response variable, competition history and experiment day as factors (including any interactions), and sample code as a random effect.

### d. Differences in community biovolume

We used the same models described above (a) but applied to the total community biovolume to estimate r and K. We fitted these models to each community replicate using biovolume data until day 20 (Phase 2, before the invasion). We then refitted these models to the entire data series (including post-invasion data) to show changes in community biovolume across the entire experiment.

### e. Community metabolic rates

We tested how competition history affected the relationship between community oxygen rates and total biovolume (Phase 2 pre-invasion), running this analysis for photosynthesis, post-illumination, and dark respiration rates separately. We also estimated daily net energy production as 12 hours of energy produced through photosynthesis minus 12 hours of respiration (calculated as 15 minutes of post-illumination and 11.75 hours of dark respiration). We excluded data from day 17 in the analysis of post-illumination rates (and thus also in the analysis of net energy production) because all samples had unusually low post-illumination rates on that day that did not fit with the trajectory observed over time.

We analysed the oxygen rates separately for the exponential (until day 10 included based on community biovolume trajectories) and stationary phase (until day 20, before invasion) because the relationship between oxygen rates and biovolume is different between these two phases (a positive relationship in exponential phase vs no relationship in stationary phase). Within each growth stage, we used linear mixed effect models with the metabolic rates as a response variable, biovolume and competition history as predictors (including their interaction), and community sample code as a random effect. Since variances were heterogeneous for the exponential phase of photosynthesis, post-illumination respiration and net energy production, we used a generalised least squares model (GLS) to apply a treatment-specific variance to each competition treatment.

### f. Community response to invader (Phase 3)

To determine how competition history influenced the response of the community (change in biomass) to the addition of an invader, we used a linear mixed effect model that includes community biovolume as a response variable (log_10_-transformed), experiment day and competition history (including their interactions) as factorial predictors, with sample code as a random effect. We fitted this model to biovolume data from day 22 (first day of invasion) to day 40 (end of experiment).

## RESULTS

### Competition history maintains species dominance but reduces evenness

Competition history initially did not affect how species performed within their community: monoculture and polyculture isolates of each species had similar maximum growth rates (r_max_) (Figure S1). As biovolume increased, however, the two weaker competitors reached a lower max. biovolume (K) in the communities of polyculture compared to the monoculture isolates (Figure 2), albeit the difference in carrying capacity (K) was significant for *Tisochrysis* (F_1,17_ = 14.12, p = 0.0016; Table S1) but not for *Dunaliella* (Figure S1, Table S1). These two weaker competitors (*Dunaliella* and *Tisochrysis*) experienced mortality (a decline in biovolume) after 14 and 8 days respectively, while the biovolume of the dominant species *Amphidinium* kept increasing. This pattern was observed in both community types. Because differences in biovolume between the dominant and the weaker competitors were more marked in the polyculture isolates (species evolved together), evenness declined faster in this treatment compared to the monoculture isolates (species evolved alone; competition history × experiment day: F_9,162_ = 4.23, p < 0.0001; Figure 3; Table S2). The dynamics of species biovolume mirrored those of cell densities because competition history did not consistently affect cell size or circularity for any of the species (Figure S2; Table S3).

**Figure 2.**
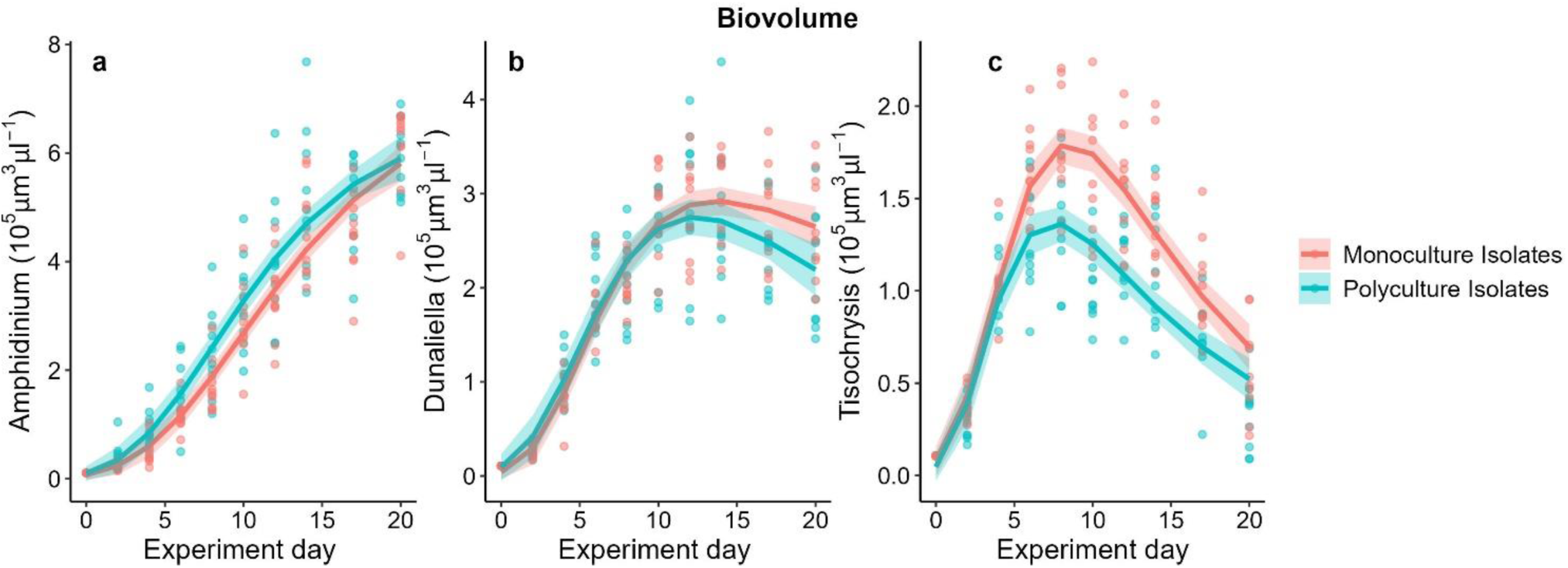
Monoculture and polyculture isolates of each species show similar biovolume growth dynamics initially, but differences in species dominance become more marked in polyculture isolates as competition increases. *Amphidinium* dominated in both treatments, whereas *Dunaliella* and *Tisochrysis* both declined from days 14 and 8 respectively – and more so in communities of polyculture isolates.

**Figure 3.**
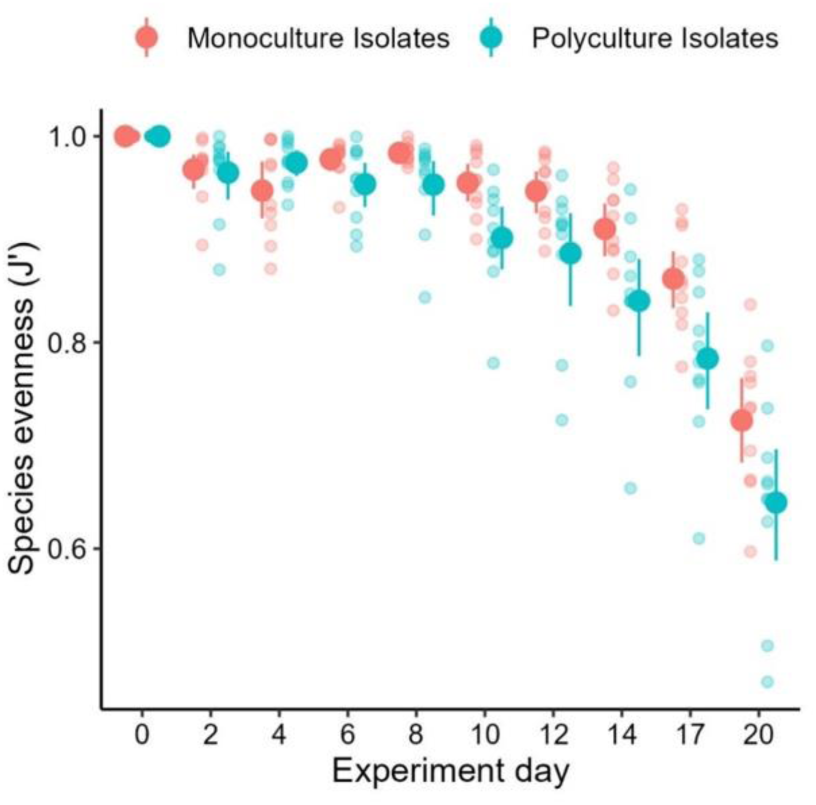
The evenness of polyculture isolates (species evolved together) declined faster over time compared to that of monoculture isolates (species evolved alone). Refer to Table S2 for statistical analysis.

### Community biovolume and oxygen production respond differently to competition history

Despite the differences in species evenness, total biovolume increased identically in the two community types (Figure 4). Monoculture and polyculture isolates had the same max. rates of increase and maximum values of total biovolume by day 20 (Figure S3, Table S4). However, the two community treatments had different oxygen evolution rates, albeit these differences were weak and mostly visible during the exponential phase.

**Figure 4.**
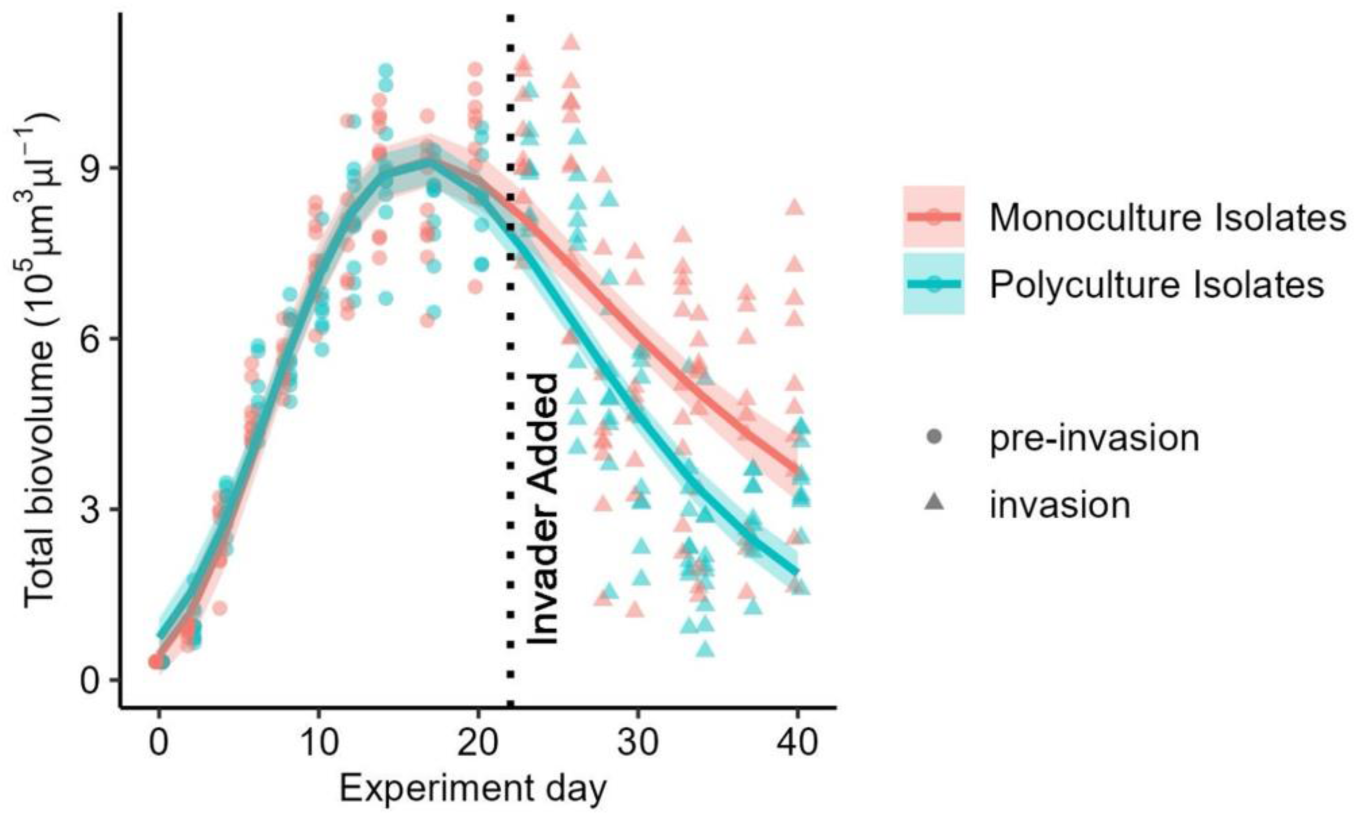
Total biovolume (μm^3^/μl) increased identically over time in communities of polyculture and monoculture isolates, up until carrying capacity (∼ day 20). The two community types then responded differently to the addition of an invader (days 22-40; triangles). The black dotted line highlights the day in which the invader (*Phaeodactylum)* was added (day 22).

The polyculture isolates tended to have higher photosynthesis (borderline competition history effect: F_1,18_ = 4.32, p = 0.0533, Table S5) and post-illumination rates (competition history effect: F_1,18_ = 7.15, p = 0.016; borderline biovolume × competition history effect: F_1,77_ = 3.94, p = 0.0508) compared to the monoculture isolates in exponential phase (Figure 5). In stationary phase, post-illumination rates were still significantly higher for the polyculture isolates (competition history: F_1,57_ = 5.6, p = 0.03), while photosynthesis rates were similar between treatments (F_1,18_ = 0.98, p = 0.34). Dark respiration rates were not affected by competition history, either in the exponential or in the stationary phase (Figure 5; Table S5). Therefore, when we estimated net energy production over 24 hours, we found the same pattern observed for photosynthesis: net energy production was marginally higher in communities of polyculture isolates during exponential phase (competition history effect: F_1,18_ = 4.37, p = 0.0496; Table S5) but not during stationary phase (F_1, 18_ = 1.92, p = 0.18).

**Figure 5.**
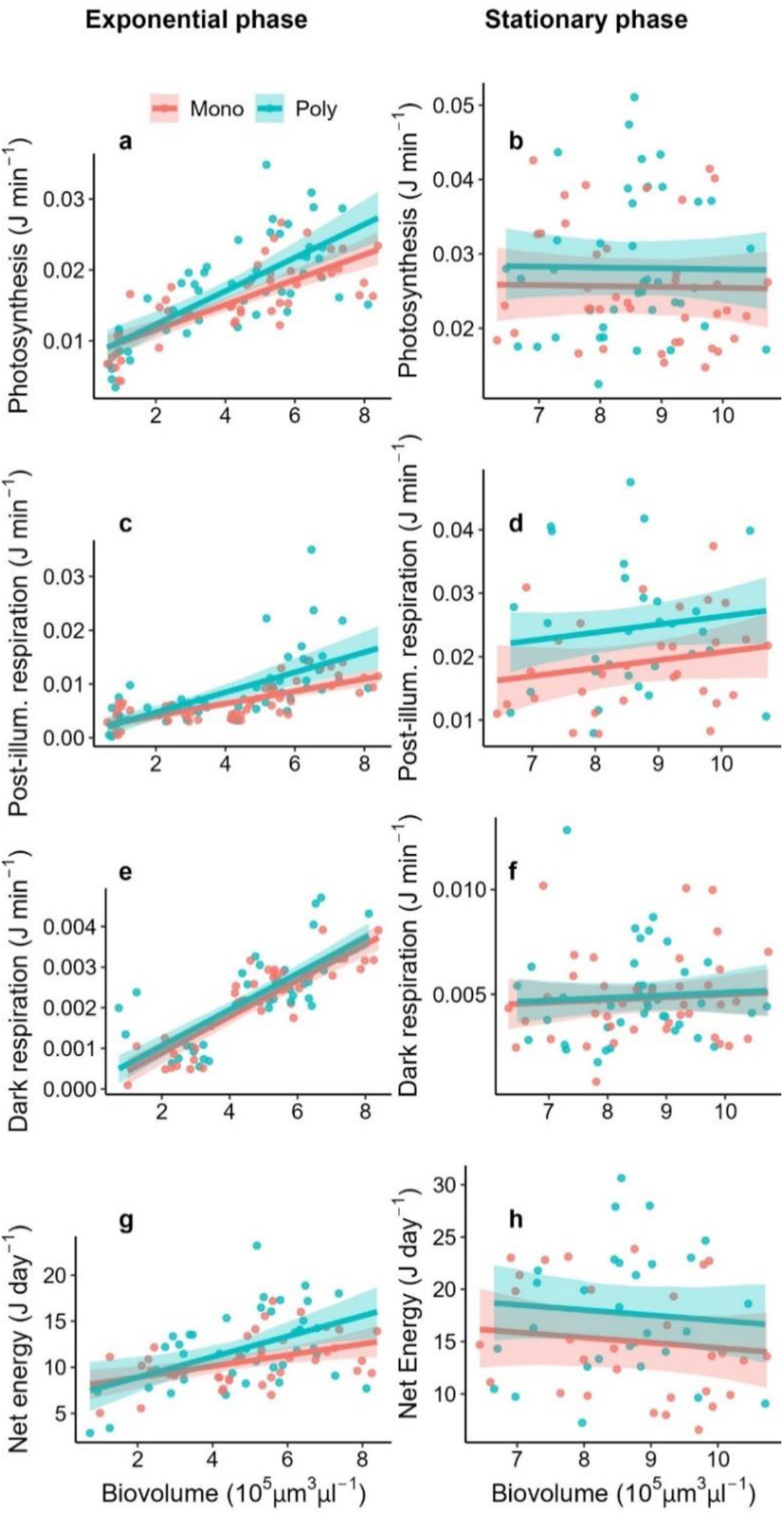
Differences in community oxygen rates as a function of total biovolume pre-invasion (up to day 20): photosynthesis (a-b), post-illumination (c-d) and dark respiration (e-f), and net daily energy production (g-h). Panels on the left refer to exponential growth phase (days 0-10), panels on the right to the stationary phase (days 12-20). Refer to Table S5 for the model outputs.

### Community response to Invasion

The addition of the invader disrupted biovolume trajectories differently depending on the competition history of the community (Figure 4). Total biovolume declined in all communities after the addition of the invader (day 22), but the polyculture isolates showed a steeper decline compared to the monoculture isolates (competition history × experiment day: F_1,138_ = 3.98, p = 0.048; Figure S4; Table S6). Despite this difference, the invader did not successfully establish in any of the cultures (except one monoculture isolate, Figure S5) and caused a decline in species biovolume in both treatments (Figure S6), which is not normally observed when cultures reach carrying capacity. The rapid loss of biovolume is particularly evident when the community trajectories are compared to those of *Amphidinium* alone (which is also the dominant species in the communities) observed in a previous experiment (Briddon *et al*. 2024), where cultures maintained a stable biovolume for several days after reaching carrying capacity (Figure S7).

## DISCUSSION

When species co-occur with competitors, they can evolve to increase niche differences and/or competitive ability. While classic theory anticipates that increased niche differences should be the most likely outcome, empirical tests more consistently show changes in competitive ability, particularly when species compete for essential resources (Germain *et al*. 2020; Hart *et al*. 2019; Miller *et al*. 2014). However, the consequences of these evolved changes for community properties are unclear. To answer this question, we investigated how evolution with multiple competitors influenced phytoplankton coexistence, productivity and stability.

Our results suggest that, even if niche partitioning is possible in phytoplankton (Stomp *et al*. 2004), evolving with multiple competitors does not significantly increase niche differentiation between these species. On the contrary, evolution with competitors weakens coexistence by strengthening competitive differences between species without altering species hierarchies. Indeed, communities of monoculture and polyculture isolates assembled in similar ways and were dominated by the same species (*Amphidinium*), but dominance patterns were stronger in polyculture isolates. These consistent patterns might arise because of trade-ups in competitive ability for multiple resources (Gallego *et al*. 2019; Passarge *et al*. 2006).

Our recent work Briddon *et al*. (2024) shows that all three species included in the communities (*Amphidinium*, *Dunaliella*, *Tisochrysis*) evolved a reduced sensitivity to intraspecific competition when tested alone, independently of whether they evolved in monoculture or polyculture. However, *Amphidinium* showed stronger changes than the other two species: it evolved the shallowest density-dependence of energy fluxes meaning that its cells produced higher amounts of net energy in dense populations (Briddon *et al*. 2024), possibly explaining its dominance in the community. *Amphidinium* is a mixotroph (Ignatiades 2012) so, when competition is intense, this species might increase energy gains by complementing photosynthesis with heterotrophy (Yang *et al*. 2021). Whatever the specific mechanism, higher rates of energy production when resources are scarce might have favored *Amphidinium* at the expense of the other species (*Tisochrysis*, *Dunaliella*).

Why is coexistence reduced in polyculture isolates? Species can evolve reduced density-dependence to cope with both intra- and inter-specific competition (Briddon *et al*. 2024; Sakarchi & Germain 2023), but evolving in a community can come at a cost (Hall *et al*. 2018; Scheuerl *et al*. 2020). Briddon *et al*. (2024) shows that changes in density-dependence were weaker in species evolved in polyculture. This reduced capacity for evolution, combined with the stronger changes in energy fluxes observed in *Amphidinium*, could explain why evenness declined faster in communities of polyculture isolates. Reduced coexistence between (co)evolved species contrasts with predictions of character displacement (Dayan & Simberloff, 2005; Pfennig & Pfennig, 2010) that however seem more likely when species compete for substitutable resources. Experiments in bacteria show that character displacement occurs (Pastore *et al*. 2021; Piccardi *et al*. 2022) and that bacteria often develop facilitative interactions via cross-feeding (Lawrence *et al*. 2012). These types of interactions however might be less common in other organisms, particularly when resources cannot be easily substituted (Zhang & Becks 2024). A limitation of our work is that, during the experimental evolution phase, species competed for nutrients but were physically separated so our results cannot account for evolutionary responses to cell-to-cell interactions. However, our results align with other studies in bacteria where evolution with multiple competitors led to maladaptation (Castledine *et al*. 2020) and slowed adaptation (Scheuerl *et al*. 2020). An important consideration might be that we evolved species under constant conditions; environmental fluctuations might be required to evolve niche differentiation (Hochfeld & Hinners 2024; Pastore *et al*. 2021). Under constant conditions, selection can drive species to convergence towards similar phenotypes. While niche convergence can be an equalising mechanism that reduces fitness differences (Chesson 2024; Leopold & Fukami 2021), it also reduces the stabilising effects that support coexistence (Lankau *et al*. 2011). Thus, in stable environments, small differences in competitive ability might over time enable a single dominant species to competitively exclude all others.

Despite differences in community evenness, there was perfect compensation with respect to biovolume at community level. The dominant species perfectly counterbalanced the decline in abundance of the competitors indicating that species loss does not necessarily affect community functioning (Gerhard *et al*. 2021). Compensatory dynamics are well documented (Dolezal *et al*. 2020; Gonzalez & Loreau 2009) and our results suggest that they might be maintained as species (co)evolve. Even as species traits and competitive ability change over time, communities can display adaptive dynamics that allow for the maintenance of community functioning (Carroll *et al*. 2023). This stability is encouraging and, even if we did not consider it here, it would be important to study compensatory dynamics as communities adapt to environmental change (Aubree *et al*. 2020).

While evolution with competitors did not alter total biovolume, it increased community oxygen fluxes, even if only transiently. It seems that competitive interactions might affect biovolume and oxygen fluxes differently, not only over ecological (Ghedini *et al*. 2022) but also evolutionary timescales, highlighting the need to measure multiple functions simultaneously (Dietrich *et al*. 2024). This result accords with the effects of eco-evolutionary dynamics observed in bacteria: Castledine *et al*. (2020) found no difference in biomass productivity between communities of monoculture- or polyculture-evolved species. While Lawrence *et al*. (2012) found that communities of polyculture isolates had significantly higher productivity (CO_2_ production rate) than communities of monoculture isolates. The differences in energy fluxes we observed in our study were marginal but still the, however little, extra energy produced by polyculture isolates was not not used for biomass production. We hypothesise that this ‘extra’ energy could have been used to generate a metabolite or cue by some of the species in the community (Zhou *et al*. 2024), such as an allelopathic cue or a toxin, that would have been produced only in polyculture (i.e. when evolved with other species) (Granato *et al*. 2019; Inglis *et al*. 2016; Lawrence *et al*. 2012). However, without analysis of the transcriptome or metabolites it is not possible to test this hypothesis.

We initially predicted that polyculture isolates might be more resistant to invasion because evolving with competitors should increase niche partitioning or resource use efficiency of the resident species (Piccardi *et al*. 2022; Spaak & Schreiber 2023). Instead, polyculture isolates were more sensitive to invasion and responded more drastically than monoculture isolates. By reducing community evenness, evolution with competitors might have affected aspects of biodiversity that are important for stability (Lepori *et al*. 2024). The invader we chose was the diatom *Phaeodactylum tricornutum*, a different functional group than the other three species but one that tends to be a strong competitor for resources (Siegel *et al*. 2020). The invader remained at low abundance in all communities, possibly because resources were nearly exhausted as communities approached stationary phase. To this respect, the communities resisted invasion because the invader did not proliferate, suggesting that communities had similar resource use patterns. Even though the invader was not able to spread, it perturbed both community types driving species towards extinction (Arnoldi *et al*. 2022; Gaertner *et al*. 2014). Competition history thus affected the speed, more than the way in which communities responded to invasion – considering whether species have a history of co(evolution) might be important for determining the rate at which their communities respond to biodiversity changes. Some of these responses might depend on whether evolution between competitors increases competitive ability more than niche differences (Lepori *et al*. 2024), so it seems important to determine in which systems evolution favours one or the other evolutionary trajectory.

## CONCLUSION

In summary, our results suggest that evolution between competitors does not majorly alter key aspects of communities, such as species hierarchies and community biomass production, but still has important consequences on their functioning. Evolving with competitors seems to reduce coexistence by strengthening competitive differences between species, at least when species compete for essential resources in stable environments. Our result is consistent with empirical and theoretical work showing that evolution is unlikely to favor coexistence (Miller *et al*. 2014; Passarge *et al*. 2006; Pastore *et al*. 2021). Importantly, small changes in community assembly and diversity can affect important aspects of community functioning, such as their oxygen production and their robustness to an invading species. If (co)evolution strengthens competitive hierarchies between resident species, communities of species that evolved together (less disturbed systems) might be more frail to invaders, which is worrying given the ongoing biodiversity changes (Loreau *et al*. 2023). Overall, these results highlight the importance of measuring multiple aspects of community functioning because the effects of multi-species evolution may not be uniform across them.

## DATA AVALIABILITY

All data and code have been deposited in Figshare and will become public upon acceptance.

## ACKNOWLEDGMENTS

GG was supported by a fellowship (LCF/BQ/PI21/11830001) from ‘‘la Caixa’’ Foundation (ID 100010434) and the European Union’s Horizon 2020 research and innovation programme under the Marie Sk1odowska Curie grant agreement no. 847648. This work was also supported by an ERC Starting Grant by the European Union to GG (ERC, META_FUN, 101116029).

## AUTHOR CONTRIBUTIONS

CB performed the research, analysed data and wrote the paper. AMG performed the research and analysed the data. GG designed the experiment, analysed data and wrote the paper.

## COMPETING INTERESTS STATEMENT

The authors declare no competing interests.

## Notes

### Competing Interest Statement

The authors have declared no competing interest.

## REFERENCES

Abrams, P.A. (1986). Character displacement and niche shift analyzed using consumer-resource models of competition. Theor Popul Biol, 29, 107–160.

Arnoldi, J.F., Barbier, M., Kelly, R., Barabás, G. & Jackson, A.L. (2022). Invasions of ecological communities: Hints of impacts in the invader’s growth rate. Methods Ecol Evol, 13, 167–182.

Aubree, F., David, P., Jarne, P., Loreau, M., Mouquet, N. & Calcagno, V. (2020). How community adaptation affects biodiversity–ecosystem functioning relationships. Ecol Lett.

Banse, K. (1991). Rates of phytoplankton cell division in the field and in iron enrichment experiments. Limnol Oceanogr, 36, 1886–1898.

Bates, D., Mächler, M., Bolker, B. & Walker, S. (2015). Fitting Linear Mixed-Effects Models Using lme4. J Stat Softw, 67.

Briddon, C.L., Estevens, R. & Ghedini, G. (2024). Evolution under competition increases phytoplankton production by reducing the density-dependence of net energy fluxes and growth.

Carroll, T., Cardou, F., Dornelas, M., Thomas, C.D. & Vellend, M. (2023). Biodiversity change under adaptive community dynamics. Glob Chang Biol, 29, 3525–3538.

Castledine, M., Padfield, D. & Buckling, A. (2020). Experimental (co)evolution in a multi-species microbial community results in local maladaptation. Ecol Lett, 23, 1673–1681.

Chesson, P. (2024). MECHANISMS OF MAINTENANCE OF SPECIES DIVERSITY.

Collins, S., Rost, B. & Rynearson, T.A. (2014). Evolutionary potential of marine phytoplankton under ocean acidification. Evol Appl, 7, 140–155.

Cooper, G.M. (2000). The Cell, A Molecular Approach. 2nd edn.

Dayan, T. & Simberloff, D. (2005). Ecological and community-wide character displacement: The next generation. Ecol Lett.

Dietrich, P., Ebeling, A., Meyer, S.T., Asato, A.E.B., Bröcher, M., Gleixner, G., et al. (2024). Plant diversity and community age stabilize ecosystem multifunctionality. Glob Chang Biol, 30.

Dolezal, J., Fibich, P., Altman, J., Leps, J., Uemura, S., Takahashi, K., et al. (2020). Determinants of ecosystem stability in a diverse temperate forest. Oikos, 129, 1692–1703.

Fox, J., W.S. (2019). An {R} Companion to Applied Regression. Third Edition. Sage.

Gaertner, M., Biggs, R., Te Beest, M., Hui, C., Molofsky, J. & Richardson, D.M. (2014). Invasive plants as drivers of regime shifts: Identifying high-priority invaders that alter feedback relationships. Divers Distrib, 20, 733–744.

Gallego, I., Venail, P. & Ibelings, B.W. (2019). Size differences predict niche and relative fitness differences between phytoplankton species but not their coexistence. ISME Journal, 13, 1133– 1143.

Gerhard, M., Mori, C. & Striebel, M. (2021). Nonrandom species loss in phytoplankton communities and its effect on ecosystem functioning. Limnol Oceanogr, 66, 779–792.

Germain, R.M., Srivastava, D. & Angert, A.L. (2020). Evolution of an inferior competitor increases resistance to biological invasion. Nat Ecol Evol, 4, 419–425.

Ghedini, G., Loreau, M. & Marshall, D.J. (2020). Community efficiency during succession: a test of MacArthur’s minimization principle in phytoplankton communities. Ecology, 101.

Ghedini, G. & Marshall, D.J. (2023). Metabolic evolution in response to interspecific competition in a eukaryote. Current Biology, 33, 2952–2961.e5.

Ghedini, G., Marshall, D.J. & Loreau, M. (2022). Phytoplankton diversity affects biomass and energy production differently during community development. Funct Ecol, 36, 446–457.

Gibbs, T., Levin, S.A. & Levine, J.M. (2022). Coexistence in diverse communities with higher-order interactions.

Gonzalez, A. & Loreau, M. (2009). The causes and consequences of compensatory dynamics in ecological communities. Annu Rev Ecol Evol Syst, 40, 393–414.

Gorter, F.A., Manhart, M. & Ackermann, M. (2020). Understanding the evolution of interspecies interactions in microbial communities. Philosophical Transactions of the Royal Society B: Biological Sciences.

Granato, E.T., Meiller-Legrand, T.A. & Foster, K.R. (2019). The Evolution and Ecology of Bacterial Warfare. Current Biology.

Guillard, R.R.L. & Ryther, J.H. (1962). STUDIES OF MARINE PLANKTONIC DIATOMS: I. CYCLOTELLA NANA HUSTEDT, AND DETONULA CONFERVACEA (CLEVE) GRAN. Can J Microbiol, 8, 229–239.

Hall, J.P.J., Harrison, E. & Brockhurst, M.A. (2018). Competitive species interactions constrain abiotic adaptation in a bacterial soil community. Evol Lett.

Hart, S.P., Turcotte, M.M. & Levine, J.M. (2019). Effects of rapid evolution on species coexistence. Proc Natl Acad Sci U S A, 116, 2112–2117.

Heggerud, C.M., Lam, K.Y. & Wang, H. (2023). Niche differentiation in the light spectrum promotes coexistence of phytoplankton species: a spatial modelling approach. J Math Biol, 86.

Hochfeld, I. & Hinners, J. (2024). Evolutionary adaptation to steady or changing environments affects competitive outcomes in marine phytoplankton. Limnol Oceanogr, 69, 1172–1186.

Ignatiades, L. (2012). Mixotrophic and heterotrophic dinoflagellates in eutrophic coastal waters of the Aegean Sea (eastern Mediterranean Sea). Botanica Marina, 55, 39–48.

Inglis, R.F., Scanlan, P. & Buckling, A. (2016). Iron availability shapes the evolution of bacteriocin resistance in Pseudomonas aeruginosa. ISME Journal, 10, 2060–2065.

Lankau, R., Jørgensen, P.S., Harris, D.J. & Sih, A. (2011). Incorporating evolutionary principles into environmental management and policy. Evol Appl, 4, 315–325.

Lawrence, D., Fiegna, F., Behrends, V., Bundy, J.G., Phillimore, A.B., Bell, T., et al. (2012). Species interactions alter evolutionary responses to a novel environment. PLoS Biol, 10.

Lear, K.O., Whitney, N.M., Morris, J.J. & Gleiss, A.C. (2021). Temporal niche partitioning as a novel mechanism promoting co-existence of sympatric predators in marine systems. Proceedings of the Royal Society B: Biological Sciences, 288.

Leopold, D.R. & Fukami, T. (2021). Greater local diversity under older species pools may arise from enhanced competitive equivalence. Ecol Lett.

Lepori, V.J., Loeuille, N. & Rohr, R.P. (2024). Robustness versus productivity during evolutionary community assembly: Short-term synergies and long-term trade-offs. Proceedings of the Royal Society B: Biological Sciences, 291.

Litchman, E., Klausmeier, C.A., Schofield, O.M. & Falkowski, P.G. (2007). The role of functional traits and trade-offs in structuring phytoplankton communities: Scaling from cellular to ecosystem level. Ecol Lett, 10, 1170–1181.

Loreau, M., Jarne, P. & Martiny, J.B.H. (2023). Opportunities to advance the synthesis of ecology and evolution. Ecol Lett, 26, S11–S15.

MacArthur, R. (1969). SPECIES PACKING, AND WHAT INTERSPECIES COMPETITION MINIMIZES*,t.

De Mazancourt, C., Johnson, E. & Barraclough, T.G. (2008). Biodiversity inhibits species’ evolutionary responses to changing environments. Ecol Lett, 11, 380–388.

De Meester, L., Brans, K.I., Govaert, L., Souffreau, C., Mukherjee, S., Vanvelk, H., et al. (2019). Analysing eco-evolutionary dynamics—The challenging complexity of the real world. Funct Ecol, 33, 43–59.

Miller, T.E., Moran, E.R. & terHorst, C.P. (2014). Rethinking niche evolution: experiments with natural communities of Protozoa in pitcher plants. Am Nat, 184, 277–283.

Oksanen J, Simpson G, Blanchet F, Kindt R, Legendre P, Minchin P, et al. (2024). vegan: Community Ecology Package.

Olito, C., White, C.R., Marshall, D.J. & Barneche, D.R. (2017). Estimating monotonic rates from biological data using local linear regression. Journal of Experimental Biology, 220, 759–764.

Padfield, D., Yvon-Durocher, G., Buckling, A., Jennings, S. & Yvon-Durocher, G. (2016). Rapid evolution of metabolic traits explains thermal adaptation in phytoplankton. Ecol Lett, 19, 133– 142.

Passarge, J., Hol, S., Escher, M. & Huisman, J. (2006). Competition for nutrients and light: Stable coexistence, alternative stable states, or competitive exclusion? Ecol Monogr, 76, 57–72.

Pastore, A.I., Barabás, G., Bimler, M.D., Mayfield, M.M. & Miller, T.E. (2021). The evolution of niche overlap and competitive differences. Nat Ecol Evol, 5, 330–337.

Pfennig, D.W. & Pfennig, K.S. (2010). Character displacement and the origins of diversity. American Naturalist, 176.

Pfennig, K.S. & Pfennig, D.W. (2009). CHARACTER DISPLACEMENT: ECOLOGICAL AND REPRODUCTIVE RESPONSES TO A COMMON EVOLUTIONARY PROBLEM. Q Rev Biol, 84, 253–276.

Piccardi, P., Alberti, G., Alexander, J.M. & Mitri, S. (2022). Microbial invasion of a toxic medium is facilitated by a resident community but inhibited as the community co-evolves. ISME Journal, 16, 2644–2652.

Pinheiro, J., B.D. (2022). nlme: linear and nonlinear mixed effects models. R package.

Pinsky, M.L. (2019). Species coexistence through competition and rapid evolution. Proc Natl Acad Sci U S A.

Pomati, F., Jokela, J., Castiglioni, S., Thomas, M.K. & Nizzetto, L. (2017). Water-borne pharmaceuticals reduce phenotypic diversity and response capacity of natural phytoplankton communities. PLoS One, 12.

Poulson-Ellestad, K.L., Jones, C.M., Roy, J., Viant, M.R., Fernández, F.M., Kubanek, J., et al. (2014). Metabolomics and proteomics reveal impacts of chemically mediated competition on marine plankton. Proc Natl Acad Sci U S A, 111, 9009–9014.

R Core Team. (2021). R: A Language and Environment for Statistical and Computing.

Reed, H.E. & Martiny, J.B.H. (2013). Microbial composition affects the functioning of estuarine sediments. ISME Journal, 7, 868–879.

Rotem, E., Loinger, A., Ronin, I., Levin-Reisman, I., Gabay, C., Shoresh, N., et al. (2010). Regulation of phenotypic variability by a threshold-based mechanism underlies bacterial persistence.

RStudio Team. (2021). RStudio: Integrated Development for R.

Sakarchi, J. & Germain, R.M. (2023). The Evolution of Competitive Ability. American Naturalist, 201, 1–15.

Scheuerl, T., Hopkins, M., Nowell, R.W., Rivett, D.W., Barraclough, T.G. & Bell, T. (2020). Bacterial adaptation is constrained in complex communities. Nat Commun, 11.

Schindelin, J., Arganda-Carreras, I., Frise, E., Kaynig, V., Longair, M., Pietzsch, T., et al. (2012). Fiji: An open-source platform for biological-image analysis. Nat Methods.

Searle, F.M.S. & Milliken, G.A. (1980). Population Marginal Means in the Linear Model: An Alternative to Least Squares Means. Am Stat, 34, 216–221.

Siegel, P., Baker, K.G., Low-Décarie, E. & Geider, R.J. (2020). High predictability of direct competition between marine diatoms under different temperatures and nutrient states. Ecol Evol, 10, 7276–7290.

Smith, B. & Wilson, J.B. (1996). A consumer’s guide to evenness indices. Oikos, 70–82.

Spaak, J.W. & Schreiber, S.J. (2023). Building modern coexistence theory from the ground up: The role of community assembly. Ecol Lett, 26, 1840–1861.

Stomp, M., Huisman, J., De Jongh, F., Veraart, A.J., Gerla, D., Rijkeboer, M., et al. (2004). Adaptive divergence in pigment composition promotes phytoplankton biodiversity. Nature, 432, 104–107.

Terhorst, C.P., Zee, P.C., Heath, K.D., Miller, T.E., Pastore, A.I., Patel, S., et al. (2018). Evolution in a community context: Trait responses to multiple species interactions. American Naturalist, 191, 368–380.

Vanvelk, H., Govaert, L., van den Berg, E.M. & De Meester, L. (2024). Eco-Evolutionary Interactions With Multiple Evolving Species Reveal Both Antagonistic and Additive Effects. Ecol Lett, 27.

Veresoglou, S.D., Xi, J. & Peñuelas, J. (2024). Mechanisms of coexistence: Exploring species sorting and character displacement in woody plants to alleviate belowground competition. Ecol Lett, 27, e14489.

White, C.R., Kearney, M.R., Matthews, P.G.D., Kooijman, S.A.L.M. & Marshall, D.J. (2011). A manipulative test of competing theories for metabolic scaling. American Naturalist, 178, 746– 754.

Wickham, H. (2011a). ggplot2: Elegant Graphics for Data Analysis. Springer.

Wickham, H. (2011b). The Split-Apply-Combine Strategy for Data Analysis. J Stat Softw, 40.

Wilke, C.O. (2016). Cowplot: Streamlined Plot Theme and Plot Annotations for ‘ggplot2.’

Williams, P.J.L.B. & Laurens, L.M.L. (2010). Microalgae as biodiesel & biomass feedstocks: Review & analysis of the biochemistry, energetics & economics. Energy Environ Sci.

Yang, H., Hu, Z. & Tang, Y.Z. (2021). Plasticity and multiplicity of trophic modes in the dinoflagellate karlodinium and their pertinence to population maintenance and bloom dynamics. J Mar Sci Eng, 9, 1–21.

Zhang, Z. & Becks, L. (2024). The mechanistic rules for species coexistence.

Zhou, Y., Wu, F., Wu, J., Overmans, S., Ye, M., Xiao, M., et al. (2024). The adaptive mechanisms of the marine diatom Thalassiosira weissflogii to long-term high CO2 and warming. Plant Journal, 119, 2001–2020.

