## Supplementary Information for "Eco-evolutionary dynamics between multiple competitors reduce phytoplankton coexistence but have limited impacts on community productivity"

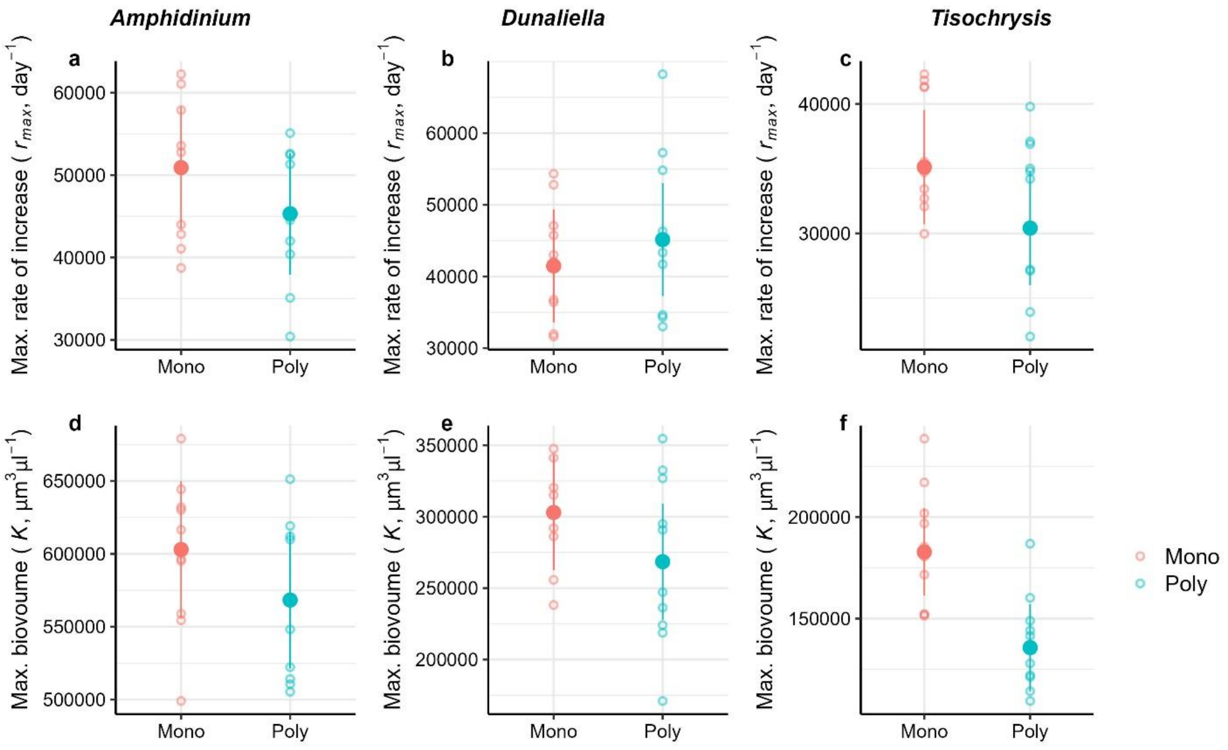

**Figure S1.** Maximum rate of increase ($r_{max}$) and max. value (K) of biovolume (μm$^3$/μl) for each species in communities of monoculture isolates (species evolved alone) or polyculture isolates (species evolved together for 4.5 months). Refer to Table S1 for the model outputs. The only significant difference was for max. biovolume for *Tisochrysis* which was higher in the monoculture isolates ($p = 0.0016$).

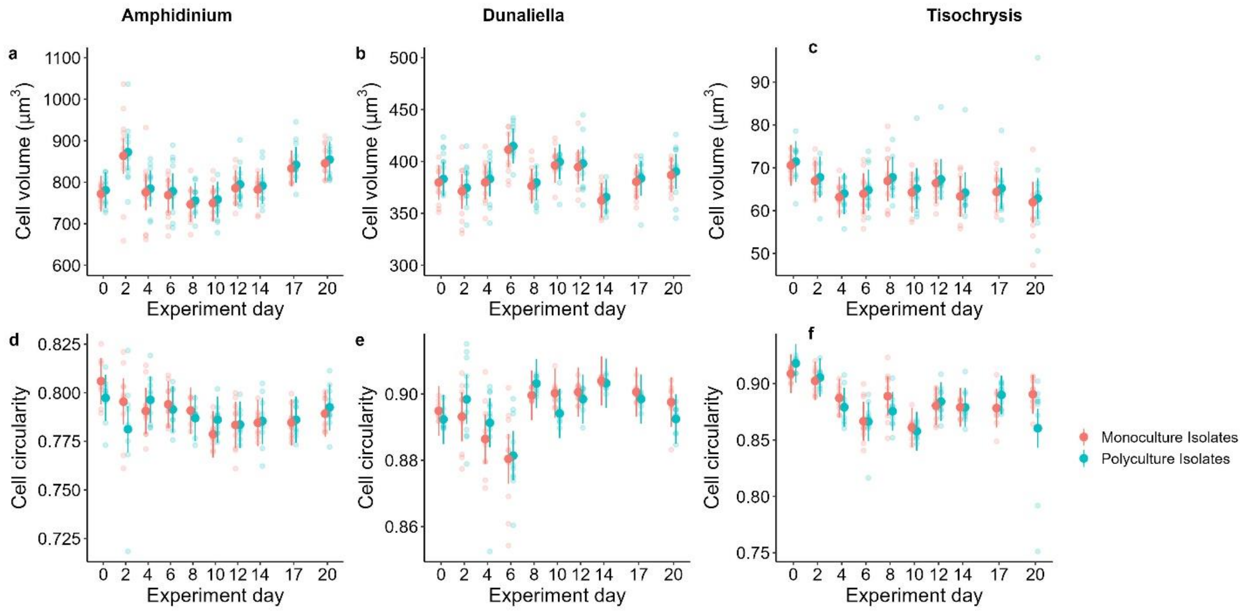

**Figure S2.** Cell size (volume) and shape (circularity) were not affected by competition history as they were similar (for each species) between monoculture and polyculture isolates. The smaller points represent the the average cell size/shape within each replicate; the larger point is the estimated marginal mean with lower and upper confidence intervals for each day of the Community Experiment (Phase 2). Refer to Table S3 for the model outputs.

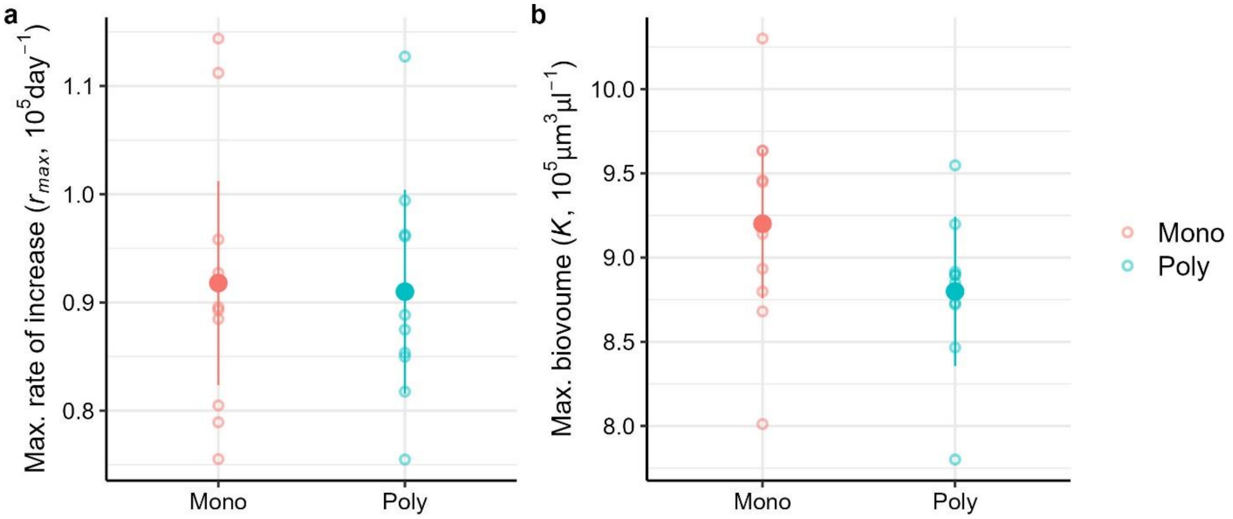

**Figure S3.** Communities of monoculture and polyculture isolates had similar maximum rate of increase ($r_{max}$) and max. value (K) of total biovolume (µm$^3$/µl). Refer to Table S4 for the model outputs.

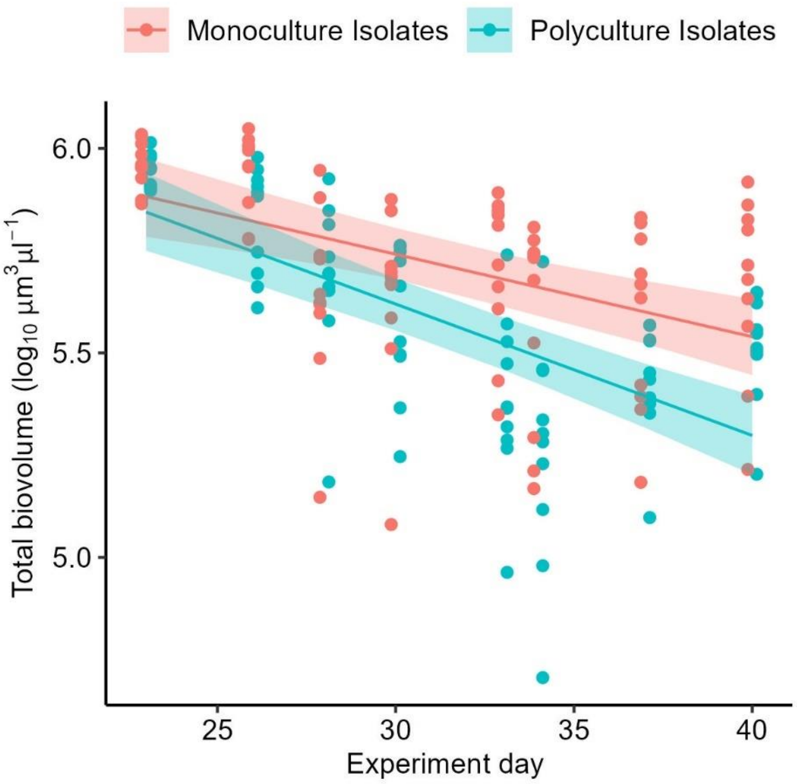

**Figure S4.** After the introduction of *Phaeodactylum* (x-axis shows experiment day, after invasion), total biovolume ($\mu m^3/\mu l$; $log_{10}$ transformed) declined faster in polyculture isolates compared to the monoculture isolates ($p = 0.048$). Refer to Table S6 for the model outputs.

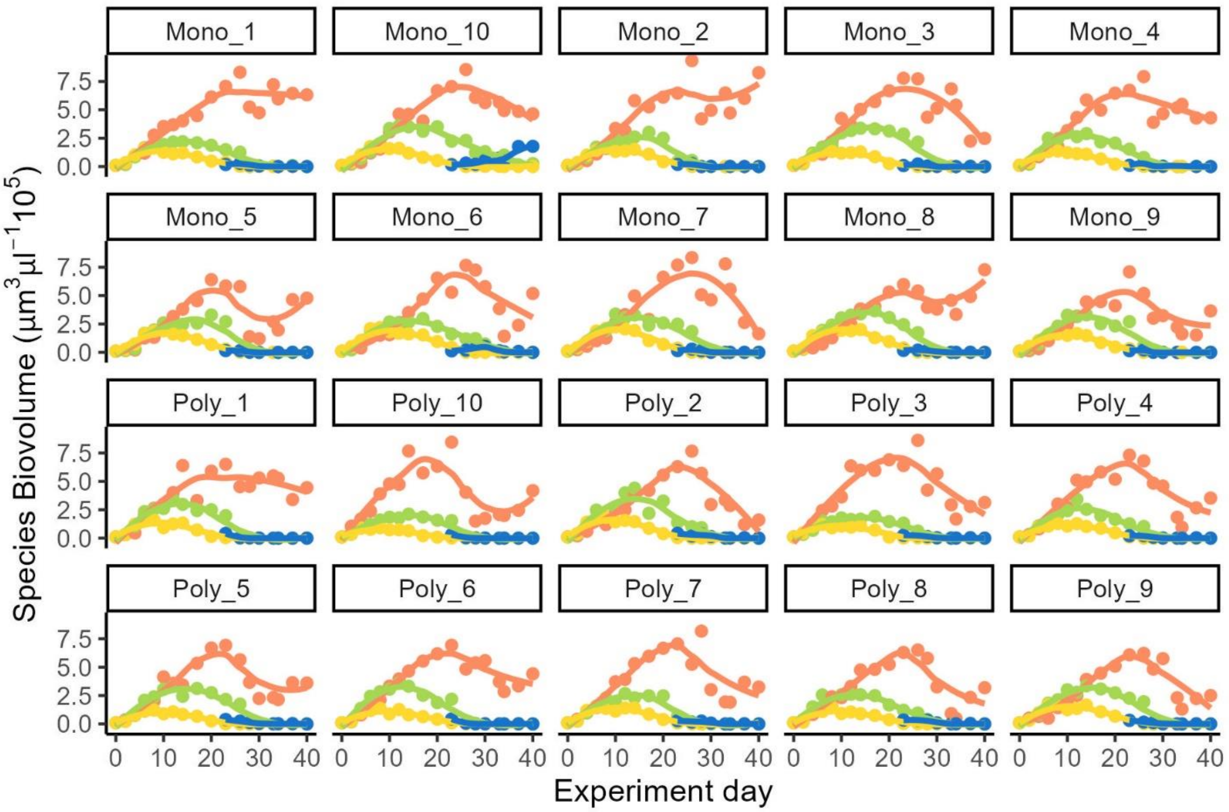

**Figure S5.** Biovolume trajectories for each species (*Amphidinium*, *Dunaliella, Tisochrysis*) in the communities of monoculture (Mono) or Polyculture (Poly) isolates, both pre- and post-invasion (marked by the addition of *Phaeodactylum* on Day 22). Data are shown for each community replicate.

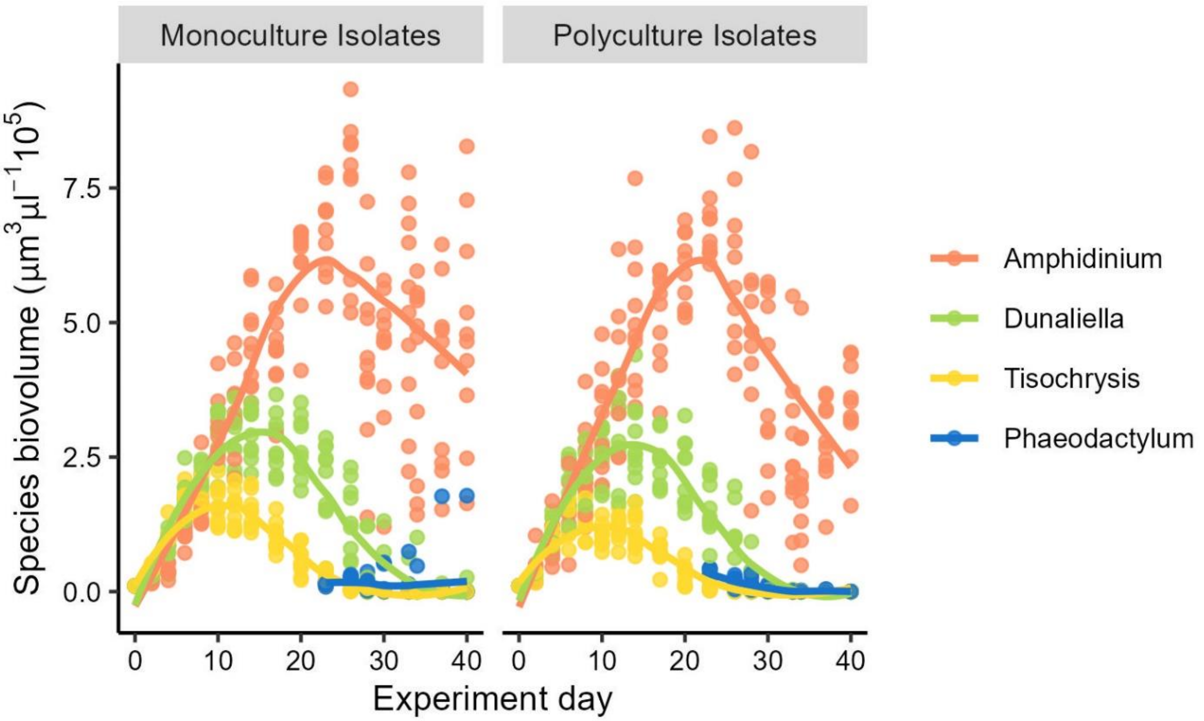

**Figure S6.** Changes in species biovolume (*Amphidinium*, *Dunaliella, Tisochrysis*), pre- and post-invasion of *Phaeodactylum* on Day 22.

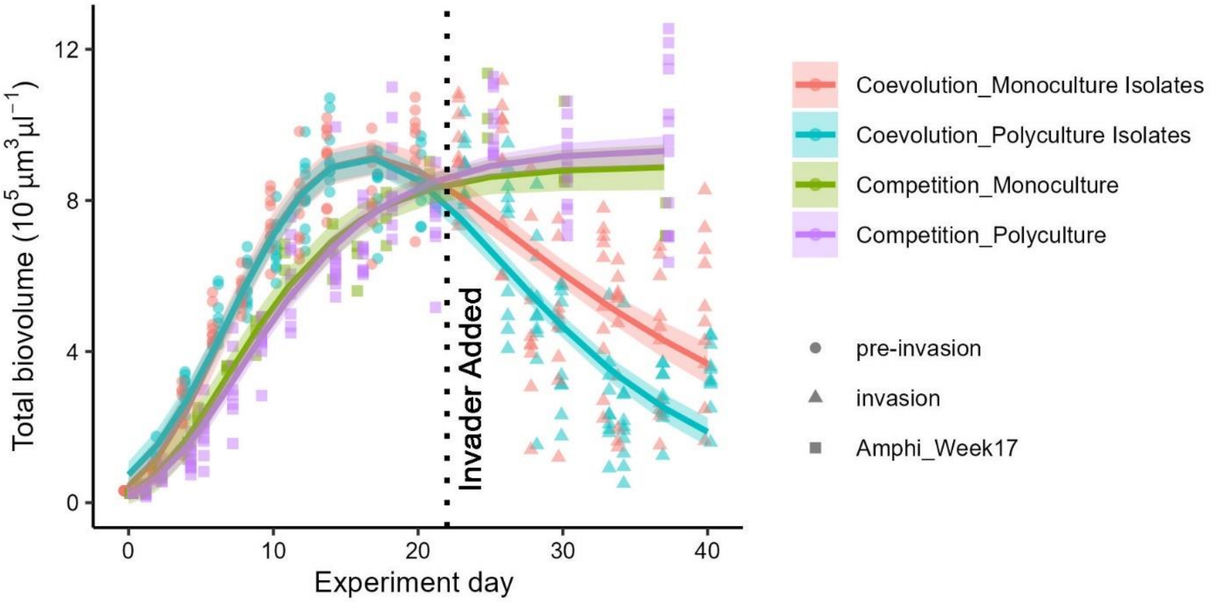

**Figure S7.** Both community treatments (polyculture and monoculture isolates) showed a rapid decline in total biovolume following invasion (from day 22). The black dotted line highlights the day in which the invader (*Phaeodactylum)* was added to the polyculture and monoculture isolates. This decline in biovolume is unexpected because culture typically remain at carrying capacity for longer periods of time, as illustrated here by the green and purple lines which show the biovolume of *Amphidinium* growing alone after 17 weeks of evolution either in monoculture or polyculture (data from Briddon *et al.* (2024)).

**Table S1**: Linear models and post-hoc test results showing the effect of competition history on the change in max. growth rates ($r_{max}$) and max. values (K) of each species biovolume ($\mu m^3/\mu l$) (*Amphidinium*, *Dunaliella*, *Tisochrysis*). Relates to Figure S1. P values < 0.05 are in bold. CL = 95% confidence level. Competition history treatments are mono = monoculture isolates, poly = polyculture isolates.

*Amphidinium* - Change in the max rates of increase

| | Df | Sum Sq | Mean Sq | F value | Pr(>F) |
|---|---|---|---|---|---|
| Biov_initial | 1 | 377289283 | 377289283 | 5.1501 | **0.03655** |
| Treatment | 1 | 105718643 | 105718643 | 1.4431 | 0.24611 |
| Residuals | 17 | 1245388665 | 73258157 | | |

*Dunaliella* - Change in the max rates of increase

| | Df | Sum Sq | Mean Sq | F value | Pr(>F) |
|---|---|---|---|---|---|
| Biov_initial | 1 | 689187732 | 689187732 | 6.75 | **0.01875** |
| Treatment | 1 | 66075471 | 66075471 | 0.6472 | 0.43224 |
| Residuals | 17 | 1735735513 | 102102089 | | |

*Tisochrysis* - Change in the max rates of increase

| | Df | Sum Sq | Mean Sq | F value | Pr(>F) |
|---|---|---|---|---|---|
| Biov_initial | 1 | 982003152 | 982003152 | 30.8096 | **<0.0001** |
| Treatment | 1 | 107069890 | 107069890 | 3.3592 | 0.0844 |
| Residuals | 17 | 541845928 | 31873290 | | |

*Amphidinium* - change in max. values of biovolume

| | Df | Sum Sq | Mean Sq | F value | Pr(>F) |
|---|---|---|---|---|---|
| Biov_initial | 1 | 1.4156e+10 | 1.4156e+10 | 4.7889 | **0.04289** |
| Treatment | 1 | 4.0581e+09 | 4.0581e+09 | 1.3728 | 0.25749 |
| Residuals | 17 | 5.0253e+10 | 2.9561e+09 | | |

*Dunaliella* - change in max. values of biovolume

| | Df | Sum Sq | Mean Sq | F value | Pr(>F) |
|---|---|---|---|---|---|
| Biov_initial | 1 | 1.1918e+09 | 1191765842 | 0.4519 | 0.5110 |
| Treatment | 1 | 6.1608e+09 | 6160793531 | 2.3360 | 0.1459 |
| Biov_init × Treatment | 1 | 5.5985e+09 | 5598501238 | 2.1228 | 0.1645 |
| Residuals | 16 | 4.2197e+10 | 2637313676 | | |

*Tisochrysis* - change in max. values of biovolume

| | Df | Sum Sq | Mean Sq | F value | Pr(>F) |
|---|---|---|---|---|---|
| Biov_initial | 1 | 7.0357e+08 | 7.0357e+08 | 0.9283 | 0.348813 |
| Treatment | 1 | 1.0704e+10 | 1.0704e+10 | 14.1224 | **0.001568** |
| Residuals | 17 | 1.2885e+10 | 7.5792e+08 | | |

| Estimated Marginal Means | Estimate | SE | Lower CL | Upper CL |
|---|---|---|---|---|
| Mono | 182813 | 8784 | 161276 | 204349 |
| Poly | 135719 | 8784 | 114183 | 157255 |

| Contrast | Estimate | SE | Df | t ratio | p value |
|---|---|---|---|---|---|
| Mono-Poly | 47094 | 12532 | 17 | 3.758 | **0.0016** |

**Table S2**: Changes in Pileou's species evenness over time between the two competition history treatments (polyculture and monoculture isolates). CL = 95% confidence level. Relates to Figure 3. P values < 0.05 are in bold.

| | NumDF | DenDF | Sum Sq | Mean Sq | F value | Pr(>F) |
|---|---|---|---|---|---|---|
| Treatment | 1 | 18 | 0.01102 | 0.010015 | 6.2685 | **0.02214** |
| Exp_day | 9 | 162 | 1.60561 | 0.178402 | 111.6610 | **<0.0001** |
| Treatment × Exp_day | 9 | 162 | 0.06086 | 0.006762 | 4.2326 | **<0.0001** |

| Estimated Marginal Means | Exp_day | Estimate | SE | Lower CL | Upper CL |
|---|---|---|---|---|---|
| Monoculture Isolates | 0 | 1 | 0.0159 | 0.968 | 1.032 |
| Polyculture Isolates | 0 | 1 | 0.0159 | 0.968 | 1.032 |
| Monoculture Isolates | 2 | 0.968 | 0.0159 | 0.936 | 0.999 |
| Polyculture Isolates | 2 | 0.965 | 0.0159 | 0.933 | 0.996 |
| Monoculture Isolates | 4 | 0.947 | 0.0159 | 0.916 | 0.979 |
| Polyculture Isolates | 4 | 0.974 | 0.0159 | 0.943 | 1.006 |
| Monoculture Isolates | 6 | 0.978 | 0.0159 | 0.946 | 1.009 |
| Polyculture Isolates | 6 | 0.954 | 0.0159 | 0.922 | 0.985 |
| Monoculture Isolates | 8 | 0.984 | 0.0159 | 0.952 | 1.015 |
| Polyculture Isolates | 8 | 0.953 | 0.0159 | 0.921 | 0.985 |
| Monoculture Isolates | 10 | 0.955 | 0.0159 | 0.923 | 0.986 |
| Polyculture Isolates | 10 | 0.901 | 0.0159 | 0.870 | 0.933 |
| Monoculture Isolates | 12 | 0.947 | 0.0159 | 0.915 | 0.979 |
| Polyculture Isolates | 12 | 0.886 | 0.0159 | 0.855 | 0.918 |
| Monoculture Isolates | 14 | 0.910 | 0.0159 | 0.878 | 0.942 |
| Polyculture Isolates | 14 | 0.840 | 0.0159 | 0.809 | 0.872 |
| Monoculture Isolates | 17 | 0.862 | 0.0159 | 0.830 | 0.894 |
| Polyculture Isolates | 17 | 0.784 | 0.0159 | 0.753 | 0.816 |
| Monoculture Isolates | 20 | 0.724 | 0.0159 | 0.693 | 0.756 |
| Polyculture Isolates | 20 | 0.645 | 0.0159 | 0.613 | 0.676 |

**Table S3**: Linear mixed effect models and post-hoc tests for changes in cell size and shape within each competition history treatment (monoculture and polyculture isolates) and experiment day (pre-invasion only, 0 to 20). Relates to Figure S2. P values < 0.05 are in bold.

| | | | | | |
|---|---|---|---|---|---|
| **Cell Size** | | | | | |
| Species = *Amphidinium* | | | | | |
| | Df | Sum Sq | Mean Sq | F value | Pr(>F) |
| Treatment | 1 | 1454 | 1454 | 0.5854 | 0.4541 |
| Exp_day | 9 | 297344 | 33038 | 13.3003 | **<0.0001** |
| Species = *Dunaliella* | | | | | |
| | Df | Sum Sq | Mean Sq | F value | Pr(>F) |
| Treatment | 1 | 239 | 239 | 0.6040 | 0.4471 |
| Exp_day | 9 | 35271 | 3919 | 9.9045 | **<0.0001** |
| Species = *Tisochrysis* | | | | | |
| | Df | Sum Sq | Mean Sq | F value | Pr(>F) |
| Treatment | 1 | 8.44 | 8.445 | 0.3509 | 0.561 |
| Exp_day | 9 | 1154.06 | 128.229 | 5.3279 | **<0.0001** |
| **Cell Shape** | | | | | |
| Species = *Amphidinium* | | | | | |
| | Df | Sum Sq | Mean Sq | F value | Pr(>F) |
| Treatment | 1 | 0.000385 | 0.000385 | 0.2864 | 0.5991 |
| Exp_day | 9 | 0.0060055 | 0.00600551 | 4.9598 | **<0.0001** |
| Treatment × Exp_day | 9 | 0.0019457 | 0.00194570 | 1.6069 | 0.1172 |
| Species = *Dunaliella* | | | | | |
| | Df | Sum Sq | Mean Sq | F value | Pr(>F) |
| Treatment | 1 | 0.0000073 | 0.00000735 | 0.1362 | 0.7164 |
| Exp_day | 9 | 0.0079192 | 0.00087992 | 16.3154 | **<0.0001** |
| Treatment × Exp_day | 9 | 0.0007130 | 0.0007922 | 1.4689 | 0.1635 |
| Species = *Tisochrysis* | | | | | |
| | Df | Sum Sq | Mean Sq | F value | Pr(>F) |
| Treatment | 1 | 0.000333 | 0.0003334 | 1.1633 | 0.295023 |
| Exp_day | 9 | 0.45281 | 0.0050312 | 17.5557 | **<0.0001** |
| Treatment × Exp_day | 9 | 0.006650 | 0.0007389 | 2.5781 | **0.008429** |

**Table S4**: Linear models and post-hoc test results showing the change in max. growth rates ($r_{max}$) and max. values (K) of total community biovolume ($\mu m^3/\mu l$) for both competition history treatments. Relates to Figure S3. P values < 0.05 are in bold. CL = 95% confidence level. Competition history treatments are mono = monoculture isolates, poly = polyculture isolates.

| Change in the max rates of increase | | | | | |
|---|---|---|---|---|---|
| | Df | Sum Sq | Mean Sq | F value | Pr(>F) |
| Biov_initial | 1 | 963519333 | 963519333 | 6.5688 | **0.02016** |
| Treatment | 1 | 3167584 | 3167584 | 0.0216 | 0.88490 |
| Residuals | 17 | 2493592519 | 146681913 | | |

| Change in the max. biovolume | | | | | |
|---|---|---|---|---|---|
| | Df | Sum Sq | Mean Sq | F value | Pr(>F) |
| Biov_initial | 1 | 2.7158e+09 | 2715845849 | 0.8383 | 0.3727 |
| Treatment | 1 | 7.9649e+09 | 7964893665 | 2.4584 | 0.1353 |
| Residuals | 17 | 5.5078e+10 | 3239879911 | | |

**Table S5**: Linear mixed effect models and the post-hoc test results on changes in community photosynthesis, post-illumination and dark respiration rates, and net energy (J/min) for the exponential and stationary phases between the two competition history treatments (polyculture vs. monoculture isolates). Related to Figure 5. CL = 95% confidence level. P values < 0.05 are in bold. Interactions were removed when p > 0.25. For rates in exponential phase (except dark respiration) we used a treatment-specific variance due to heterogeneous variances. *borderline significance.

**Photosynthesis – Exponential Phase**

| | NumDF | DenDF | F value | P value |
|---|---|---|---|---|
| Intercept | 1 | 77 | 1109.8127 | **<0.0001** |
| Biovolume | 1 | 77 | 128.9058 | **<0.0001** |
| Treatment | 1 | 18 | 4.3170 | 0.0523 * |
| Biovolume × Treatment | 1 | 77 | 2.4458 | 0.1219 |

Posthoc test for treatment

| Estimated Marginal Means | Estimate | SE | Lower CL | Upper CL | |
|---|---|---|---|---|---|
| Monoculture Isolates | 0.0155 | 0.000634 | 0.0141 | 0.0168 | |
| Polyculture Isolates | 0.0175 | 0.000770 | 0.0159 | 0.0191 | |
| Contrast | Estimate | SE | Df | t ratio | p value |
| Monoculture Isolates - Polyculture Isolates | -0.00206 | 0.000997 | 18 | -2.069 | 0.0533 * |

**Photosynthesis – Stationary Phase**

| | NumDF | DenDF | Sum Sq | Mean Sq | F value | Pr(>F) |
|---|---|---|---|---|---|---|
| Biovolume | 1 | 69.759 | 1.3280e-06 | 1.3280e-06 | 0.0206 | 0.8864 |
| Treatment | 1 | 17.974 | 6.3128e-05 | 6.3128e-05 | 0.9778 | 0.3359 |

**Post-Illumination Respiration – Exponential Phase**

| | NumDF | DenDF | F value | P value |
|---|---|---|---|---|
| Intercept | 1 | 77 | 408.2274 | **<0.0001** |
| Biovolume | 1 | 77 | 93.0053 | **<0.0001** |
| Treatment | 1 | 18 | 7.1518 | **0.0155** |
| Biovolume × Treatment | 1 | 77 | 3.9362 | 0.0508 * |

Posthoc test for biovolume × treatment effect

| Estimated Marginal Means | Estimate | SE | Lower CL | Upper CL | |
|---|---|---|---|---|---|
| Monoculture Isolates | 0.00114 | 0.000141 | 0.000861 | 0.00142 | |
| Polyculture Isolates | 0.00187 | 0.000336 | 0.001196 | 0.00253 | |
| Contrast | Estimate | SE | Df | t ratio | p value |
| Monoculture Isolates - Polyculture Isolates | -0.000723 | 0.000365 | 77 | -1.984 | 0.0508 * |

**Post-Illumination Respiration – Stationary Phase**

| | NumDF | DenDF | Sum Sq | Mean Sq | F value | Pr(>F) |
|---|---|---|---|---|---|---|
| Biovolume | 57 | 1 | 0.00010863 | 0.00010863 | 1.3164 | 0.25603 |
| Treatment | 57 | 1 | 0.00046207 | 0.00046207 | 5.5997 | **0.02139** |

Posthoc test for treatment only effect

| Estimated Marginal Means | Estimate | SE | Lower CL | Upper CL | |
|---|---|---|---|---|---|
| Monoculture Isolates | 0.0189 | 0.00166 | 0.0154 | 0.0224 | |
| Polyculture Isolates | 0.0245 | 0.00166 | 0.0210 | 0.0280 | |
| Contrast | Estimate | SE | Df | t ratio | p value |
| Monoculture Isolates - Polyculture Isolates | -0.00557 | 0.00235 | 17.6 | -2.366 | **0.0297** |

**Dark Respiration – Exponential Phase**

| | NumDF | NumDF | Sum Sq | Mean Sq | F value | Pr(>F) |
|---|---|---|---|---|---|---|
| Biovolume | 1 | DenDF | 5.7677e-05 | 5.7677e-05 | 150.7939 | **<0.0001** |
| Treatment | 1 | 80 | 5.9700e-07 | 5.9700e-07 | 1.5614 | 0.2151 |

**Dark Respiration – Stationary Phase**

| | NumDF | DenDF | Sum Sq | Mean Sq | F value | Pr(>F) |
|---|---|---|---|---|---|---|
| Biovolume | 1 | 76 | 1.5602e-06 | 1.5602e-06 | 0.3299 | 0.5674 |
| Treatment | 1 | 76 | 1.5744e-07 | 1.5744e-07 | 0.0333 | 0.8557 |

**Net Energy – Exponential Phase**

| | NumDF | DenDF | F value | P value |
|---|---|---|---|---|
| Intercept | 1 | 61 | 1197.4274 | **<0.0001** |
| Biovolume | 1 | 61 | 19.5866 | **<0.0001** |
| Treatment | 1 | 18 | 4.3725 | 0.0510 * |
| Biovolume × Treatment | 1 | 61 | 1.9001 | 0.1731 |

**Posthoc test for treatment only effect**

| Estimated Marginal Means | Estimate | SE | Lower CL | Upper CL |
|---|---|---|---|---|
| Monoculture Isolates | 10.6 | 0.388 | 9.82 | 11.4 |
| Polyculture Isolates | 12.1 | 0.569 | 10.89 | 13.3 |

| Contrast | Estimate | SE | Df | t ratio | p value |
|---|---|---|---|---|---|
| Monoculture Isolates - Polyculture Isolates | -1.45 | 0.689 | 18 | -2.105 | **0.0496** |

**Net Energy – Stationary Phase**

| | NumDF | DenDF | Sum Sq | Mean Sq | F value | Pr(>F) |
|---|---|---|---|---|---|---|
| Biovolume | 1 | 51.319 | 14.861 | 14.861 | 0.5748 | 0.4518 |
| Treatment | 1 | 18.290 | 49.622 | 49.622 | 1.9192 | 0.1826 |

**Table S6**: Linear mixed effect models and post-hoc test results showing the effect of the invasion of *Phaeodactylum* (day 21 onwards) on total biovolume ($\mu m^3/\mu l$) for both competition history treatments. Relates to Figure S4. P values < 0.05 are in bold. CL = 95% confidence level.

| | NumDF | DenDF | Sum Sq | Mean Sq | F value | Pr(>F) |
|---|---|---|---|---|---|---|
| Exp_day | 1 | 138 | 3.06600 | 3.06600 | 74.8809 | **<0.0001** |
| Treatment | 1 | 150.88 | 0.06399 | 0.06399 | 1.5629 | 0.21318 |
| Exp_day × Treatment | 1 | 138 | 0.16299 | 0.16299 | 3.9807 | **0.04799** |
| Posthoc test for exp day × treatment effect | | | | | | |
| Estimated Marginal Means | Estimate | | SE | Lower CL | Upper CL | |
| Monoculture Isolates | -0.0200 | | 0.00424 | -0.0283 | -0.0116 | |
| Polyculture Isolates | -0.0319 | | 0.00424 | -0.0403 | -0.0235 | |
| Contrast | Estimate | | SE | Df | t ratio | p value |
| Monoculture Isolates-Polyculture Isolates | 0.012 | | 0.00599 | 138 | 1.995 | **0.048** |
